# Long non-coding RNAs regulate the expression of cell surface receptors in plants

**DOI:** 10.1101/2024.04.22.590565

**Authors:** Hemal Bhasin, Hasna Khan, Zachary Kileeg, G. Adam Mott

## Abstract

Plants are exposed to a variety of growth, developmental, and environmental cues during their lifespan. To survive and thrive, plants have developed sophisticated ways of responding to these signals that involve regulation at the transcriptional, post-transcriptional, translational, and post-translational levels. Leucine-rich repeat receptor-like kinases are the largest family of receptor-like kinases in plants and respond to a range of external and internal stimuli. They act as crucial regulators of plant growth, development, and immunity. To fully understand LRR-RLK function, it is essential to understand how their expression is regulated under different conditions. While there have been numerous studies on post-translational regulation of LRR-RLKs through phosphorylation and ubiquitination, there is little known about the mechanisms of transcriptional and post-transcriptional regulation of LRR-RLKs. In this study, we show that natural antisense transcript long non-coding RNAs are central regulators of LRR-RLK expression at the transcriptional and post-transcriptional levels. LRR-RLK genes are almost universally associated with cis-NATs and we confirm cis-NAT expression *in planta* using strand-specific RT-PCR. We leverage several well-studied LRR-RLKs to demonstrate that cis-NATs regulate LRR-RLK expression and function. For cis-NATs to fine-tune LRR-RLK expression, their expression and regulatory activity must be tightly controlled and cell autonomous. Using a combination of GUS reporter assays and tissue-specific promoters, we provide evidence that cis-NATs have these characteristics, positioning them as key regulators of LRR-RLK function. We also demonstrate that the association of LRR-RLK genes with cis-NATs is conserved across much of plant evolution, suggesting that this previously unexplored regulatory mechanism serves an important and ancient purpose.

## Introduction

Plants are constantly exposed to a diverse range of external and internal stimuli. These incoming signals must be integrated to guide an appropriate adaptive response and optimize plant fitness. This process requires extremely complex gene regulation involving epigenetic, transcriptional, post-transcriptional, and translational control. Strand-specific transcriptomics has recently identified long non-coding RNAs (lncRNAs) as an important class of gene expression regulators that control diverse developmental and environmental responses in plants(1–3).

An emerging class of lncRNAs are natural antisense transcript lncRNAs (NAT lncRNAs, hereafter NATs) that are transcribed from the antisense strand of coding genes and are widespread in plants and animals(4–8). NATs are grouped into two categories, cis-NATs and trans-NATs, based on their locus of action(4, 9). Trans-NATs act at a distal site from their genomic location and possess only partial sequence complementarity to the regulated sense transcript(10). Cis-NATs act on the sense strand of the same genomic locus and therefore have perfect sequence complementarity to the genes they regulate. Large-scale sequencing efforts have identified cis-NATs associated with 15-20% of *Arabidopsis thaliana* (hereafter Arabidopsis) genes(7, 11, 12).

NATs can act as positive or negative regulators of their target genes via various mechanisms acting at the transcriptional, post-transcriptional, or translational level(7, 13–16). Some cis-NATs also serve as a source of small interfering RNAs (NAT-siRNAs) that play known roles in environmental and developmental responses, though recent studies show that this mechanism of regulation is rare(7, 15). NAT expression can be inducible and is often specific to a developmental stage, tissue, or stress condition, enabling them to fine-tune the expression of their targets(7). Many genome-wide studies have surveyed cis-NAT expression in response to different developmental and environmental cues, but since the effect and mechanism of regulation can only be determined experimentally, very few cis-NATs have been characterized in detail(11, 12, 15, 17). The known examples are involved in the control of developmental and abiotic stress responses such as seed germination, flowering time control, adaptation to cold, cytokinin production, and nutrient response(13, 15, 18–21).

Plants must be able to precisely detect diverse stimuli and transduce them into intracellular signalling patterns to accurately regulate gene expression. To detect extracellular stimuli, Arabidopsis relies heavily on its genomic repertoire of ∼600 receptor-like kinases (RLKs)(22). The largest family of RLKs in plants is the leucine-rich repeat receptor-like kinases (LRR-RLKs), with ∼225 members in Arabidopsis(22, 23). While the majority of LRR-RLKs have no known biological function, the characterized examples regulate growth, development, hormone signalling, and immunity(24). Two well-studied LRR-RLKs involved in plant growth and development are BRASSINOSTEROID INSENSITIVE 1 (BRI1), which is the receptor for the brassinosteroid family of growth-promoting steroid hormones in Arabidopsis(25), and CLAVATA1 (CLV1), which is the receptor for the CLV3 peptide and plays an important role in shoot meristem maintenance(26). LRR-RLKs also play key roles in plant immunity through the perception of microbe-associated molecular patterns (MAMPs) and initiation of intracellular immune signalling(27). Some LRR-RLKs also regulate plant immunity by acting as signalling competent co-receptors for other LRR-RLKs or related receptor-like protein (RLPs). SUPPRESSOR OF BIR1-1 (SOBIR1) is an important LRR-RLK that plays such a role via its interactions with numerous RLPs and is required for their immune regulatory function(28, 29).

Several lines of evidence show that LRR-RLK expression is highly regulated, including in response to stimuli such as MAMP perception(30–35). As LRR-RLKs physically interact to regulate function and form a unified regulatory network(36, 37), the regulation of any single LRR-RLK affects the entire signaling network. Therefore, to fully understand how LRR-RLKs control important developmental and immune processes as a unified network, we must determine how LRR-RLK family expression and activation are fine-tuned under different stimuli. LRR-RLKs may be regulated at the level of transcription, post-transcription, translation, or post-translation. While there have been numerous studies on post-translational regulation of LRR-RLKs through phosphorylation and ubiquitination, there is little known about the mechanisms of transcriptional and post-transcriptional control of LRR-RLKs.

In this study, we identify cis-NATs as key regulators of LRR-RLK expression and function. We experimentally confirm that the majority of LRR-RLKs are associated with expressed cis-NATs. Using well-studied LRR-RLKs as examples, we demonstrate that the associated cis-NATs can regulate LRR-RLK expression and function. We further show that the promoters of these cis-NATs are active and show distinct expression patterns *in planta* and confirm the cell-specificity of this regulation by deploying tissue layer-specific promoters to target LRR-RLK function in relevant tissues. Finally, we demonstrate that the association between LRR-RLKs and cis-NATs is not limited to Arabidopsis, but is conserved in diverse, agronomically important crops. Our findings reveal a critical mechanism for the global fine-tuning of LRR-RLK expression and function.

## Results

### LRR-RLKs are globally associated with cis-NATs

Cis-NATs exhibit dynamic regulation with distinct expression patterns and therefore have the potential to fine-tune gene regulation in response to stimuli. Given these characteristics, we hypothesized that cis-NATs may provide dynamic control of LRR-RLKs. To test this hypothesis, we identified cis-NATs associated with LRR-RLK genes by mining publicly available strand-specific RNA sequencing and directional tiling array datasets (Figure 1A). The pool of candidate cis-NAT sequences was taken from three sources: a total of 4,411 novel cis-NATs identified by strand-specific sequencing from Arabidopsis seedlings grown under various conditions by Deforges et al.(12), a total of 37,238 NAT pairs identified in Arabidopsis using the Reproducibility-based Tiling-array Analysis Strategy on 200 directional tiling array datasets by Wang et al.(11, 38), and the annotated cis-NATs found in Araport11(39, 40). Comparing these predicted cis-NATs to the location of known LRR-RLK genes, we identified a total of 70 cis-NATs associated with LRR-RLK genes from Deforges et al., with a further 200 from Wang et al., and 20 from Araport11. We excluded mRNAs from pairs of protein-coding genes that overlap in opposite orientations and removed duplicates found in multiple sources. This analysis resulted in a candidate pool of 212 cis-NATs uniquely associated with an LRR-RLK gene (Table S1, Figure 1B).

**Figure 1.**
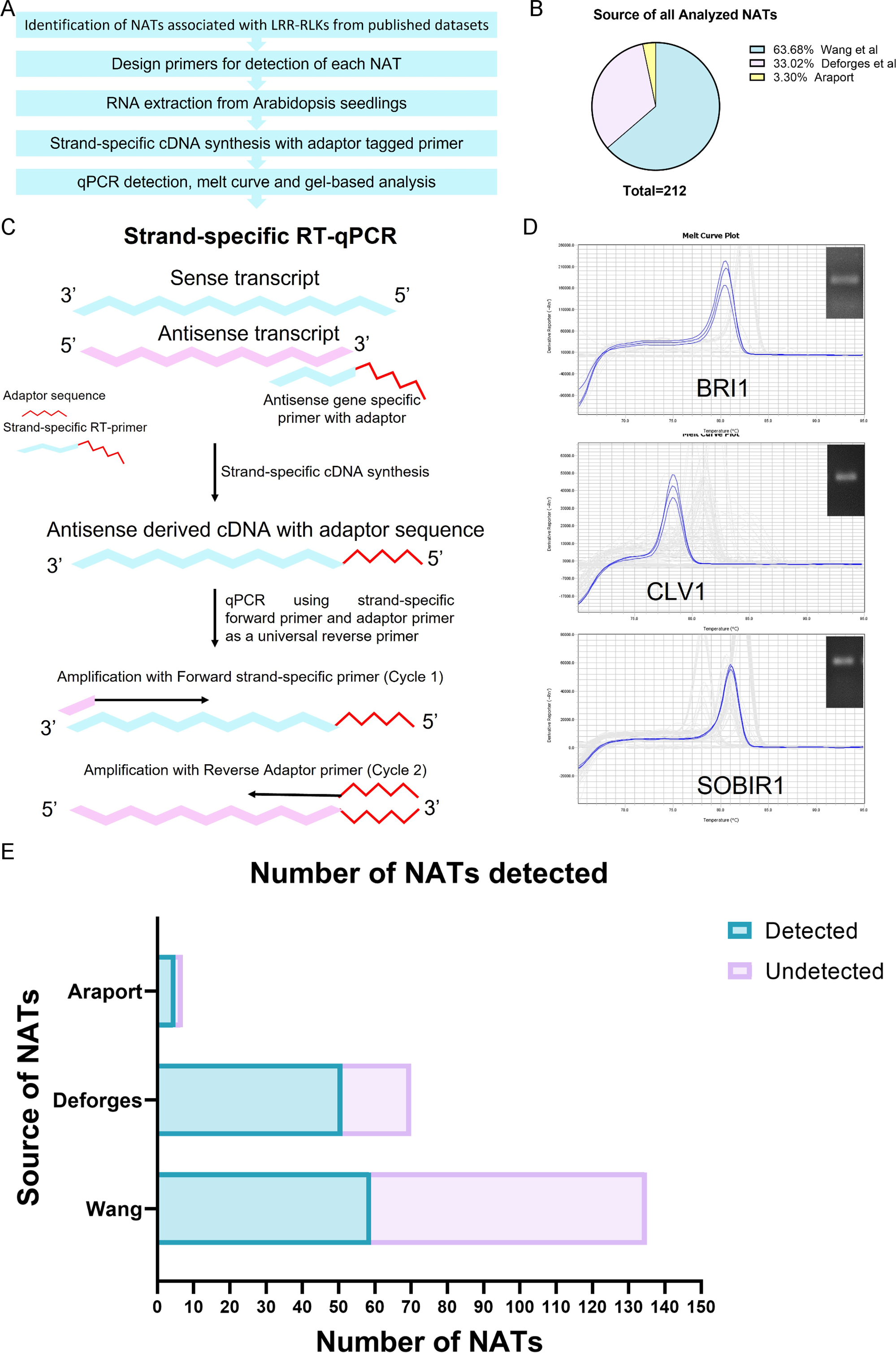
Screening for cis-NATs associated with LRR-RLKs. (A) Schematic of steps involved in the detection of LRR-RLK associated cis-NATs. (B) Source of all the cis-NATs tested. (C) Diagram explaining strand-specific qRT-PCR with a tagged primer. (D) Example of qPCR detection and gel analysis of *BRI1_NAT*, *CLV1_NAT* and *SOBIR1_NAT*. Image shows the representative qPCR melt curve displaying a single strong peak, inset is the product as seen visualized following agarose gel electrophoresis. (E) Statistics showing total number of LRR-RLK associated cis-NATs analysed and detected.

The expression of all 212 cis-NATs were validated in 10-day-old Arabidopsis seedlings using a highly sensitive strand-specific RT-PCR assay(19, 41) (Figure 1A, C). Product specificity was confirmed for each cis-NAT using melt curve and agarose gel analysis (Figure 1D, Figure S1). Based on this analysis, we could confirm *in planta* expression of 51 of 70 (72.8%) cis-NATs from Deforges et al. 2019, 59 of 135 (43.7%) cis-NATs from Wang et al. 2014, and 5 of 7 (71.4%) cis-NATs from Araport11 in tissue from this single experimental condition (Figure 1E).

### Overexpression of cis-NATs leads to functional phenotypes

Having identified cis-NATs overlapping with the vast majority of LRR-RLK genes and validated their expression *in planta*, we sought to test if cis-NAT expression alters phenotypes. We selected three candidate cis-NATs for detailed analysis: the cis-NATs associated with *BRI1 (BRASSINOSTEROID INSENSITIVE 1)*, *CLV1 (CLAVATA 1)*, and *SOBIR1 (SUPPRESSOR OF BIR1-1)*, hereafter *BRI1_NAT*, *CLV1_NAT* and *SOBIR1_NAT* respectively (Figure 1C). These candidates were chosen because they were abundantly expressed and overlapped LRR-RLKs known to have critical roles in plant development or immunity(28, 42–47) (Figure S2). We overexpressed each of these cis-NATs in Arabidopsis using the constitutive 35S promoter from the Cauliflower Mosaic Virus (CaMV)(48) and analyzed transgenic plants to confirm increased transcript levels and identify any obvious phenotypes. While the cis-NATs are named for their genomic relationship to the proposed locus of action, several studies have shown that regulation can occur when the NAT is expressed in trans(12, 15, 19).

The overexpression of *BRI1_NAT* resulted in the majority of T1 lines displaying stunted, dwarf phenotypes with a leaf morphology that is reminiscent of the *bri1* knockout phenotype(43) (Figure 2A, B). A total of 97 transgenic plants were analysed in T1, with 94 showing this phenotype. Of these, 72 exhibited strong *bri1*-mutant-like phenotypes that resulted in mostly sterile plants, and 22 exhibited more moderate *bri1*-mutant-like phenotypes with limited fertility. This suggests that the *BRI1_NAT* antagonizes *BRI1* function.

**Figure 2.**
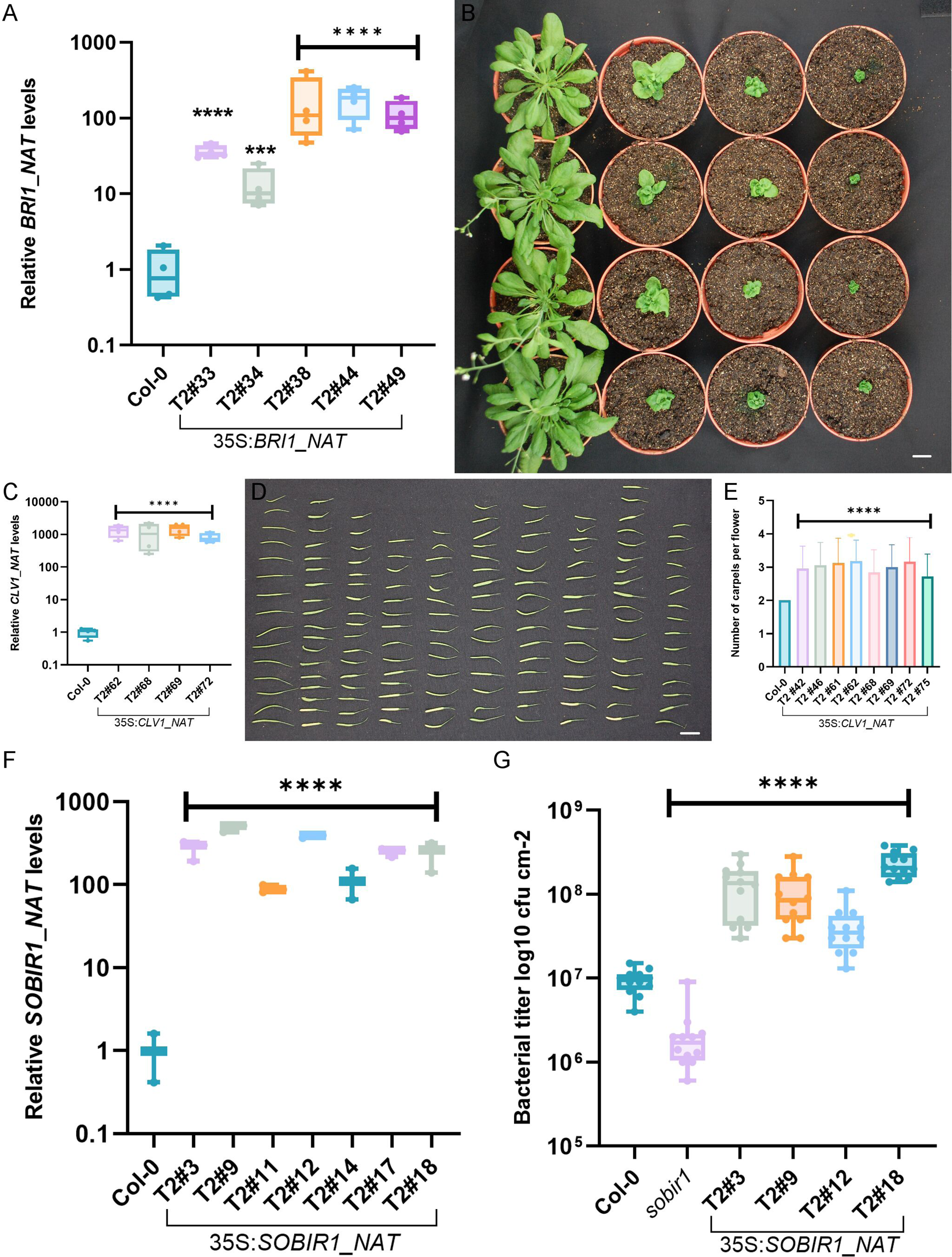
Overexpression of cis-NATs leads to functional phenotypes in transgenic lines. (A) Levels of *BRI1_NAT* in transgenic lines overexpressing *BRI1_NAT*. For all qPCR analyses, data was normalized using two internal controls – *Ubiquitin 2* and *PP2A*. For ease of visualization, the mean value of all Col-0 biological replicates was set to 1 and every data point was then calculated as fold-change relative to that value. Significance is determined by one-way ANOVA followed by Dunnett’s test for sample comparisons with Col-0 as control. Four biological samples and three technical replicates were analyzed, with the experiment repeated three times. (**** = p < 0.0001, ***= p< 0.001). Closed circles represent the mean value for individual biological replicates, the boxes represent the first and third quartiles, the horizontal line indicates the median, and whiskers extend to the farthest points not considered outliers. (B) Phenotype of plants overexpressing *BRI1_NAT* under 35S promoter. Starting from left, the first column contains 7-week-old Col-0 plants, the following three columns each consist of four independent *BRI1_NAT* overexpression lines, with phenotypic severity increasing from left to right (plants display a representative phenotype observed across several independent T1 lines). Scale bar = 1 cm. (C) Levels of *CLV1_NAT* transgenic lines overexpressing *CLV1_NAT* under 35S promoter. Significance is determined by one-way ANOVA followed by Dunnett’s test for sample comparisons with Col-0 as control. Four biological samples and three technical replicates were analyzed, with the experiment repeated three times. (**** = p < 0.0001). (D) Silique phenotype of plants overexpressing *CLV1_NAT*. Starting from the left, the first column contains siliques from Col-0 plants, the second column contains siliques from *clv-101* knockout mutant, and the remaining columns show siliques from independent *CLV1_NAT* overexpressing transgenic lines. Scale bar = 1 cm. (E) Carpel counts in plants overexpressing *CLV1_NAT* under 35S promoter. Significance is determined by one-way ANOVA followed by Dunnett’s test for sample comparisons with Col-0 as control. Ten siliques from each of five independent plants were examined for each genotype. (**** = p < 0.0001). (F) Levels of *SOBIR1_NAT* in transgenic lines overexpressing *SOBIR1_NAT*. Significance is determined by one-way ANOVA followed by Dunnett’s test for sample comparisons with Col-0 as control. Three biological samples and three technical replicates were analyzed, with the experiment repeated three times. (**** = p < 0.0001). (F) Growth of *Pseudomonas syringae* pv. *tomato* strain DC3000 (*Pst*) in 5-week-old *Arabidopsis thaliana* plants of the indicated genotypes. DC3000 titers were evaluated at 3 days post-infection. For each genotype, 12 individual plants (with 4 leaf discs from each plant) were harvested 3 days after infiltration and the experiment was repeated thrice with similar results. Significance is determined by one-way ANOVA followed by Dunnett’s test (two-tailed) for sample comparisons with Col-0 as control. (**** = p < 0.0001).

The overexpression of *CLV1_NAT* resulted in transgenic plants displaying club-shaped siliques and increased carpel number, a phenotype similar to the one observed when CLV1 function is disrupted (46, 47, 49) (Figure 2C, D, E). A total of 39 T1 transgenic plants were analyzed, with 20 displaying such abnormalities, with phenotypic severity ranging from strong to moderate. This suggests that the *CLV1_NAT* antagonizes *CLV1* function.

*SOBIR1_NAT* overexpression did not produce any obvious morphological phenotype, despite several transgenic lines showing elevated *SOBIR1_NAT* expression (Figure 2F). We tested the immune response of these transgenic lines, as SOBIR1 plays a crucial role in plant immune signalling(28). When challenged with pathogenic *Pseudomonas syringae pv.* tomato *(Pst)* DC3000, *SOBIR1_NAT* overexpressing lines consistently showed enhanced bacterial susceptibility (Figure 2G). A T-DNA knockout mutant of SOBIR1 showed the opposite phenotype with reduced bacterial growth compared to Col-0 (Figure 2G). This suggests that the *SOBIR1_NAT* is a functional agonist of *SOBIR1*.

These results show that cis-NAT changing the dosage of LRR-RLK-associated cis-NATs in Arabidopsis has functional consequences for the cognate LRR-RLK.

### Overexpression of cis-NATs regulates LRR-RLK levels in transgenic lines

To test if cis-NAT over-expression phenotypes were a result of modifying the expression of cognate LRR-RLKs, we measured LRR-RLK expression in cis-NAT overexpression lines using qPCR, ensuring that the experimental design specifically detects the LRR-RLK mRNA rather than the cis-NAT (details in Materials and Methods). We observed a significant reduction in mRNA levels of *BRI1* and *CLV1* in the *BRI1_NAT* and *CLV1_NAT* overexpression lines respectively, which is consistent with the *bri1* and *clv1* mutant-like phenotypes observed in these lines (Figure 3A, B). *SOBIR1* mRNA levels in *SOBIR_NAT* overexpression lines were not significantly different from those in Col-0 plants (Figure 3C).

**Figure 3.**
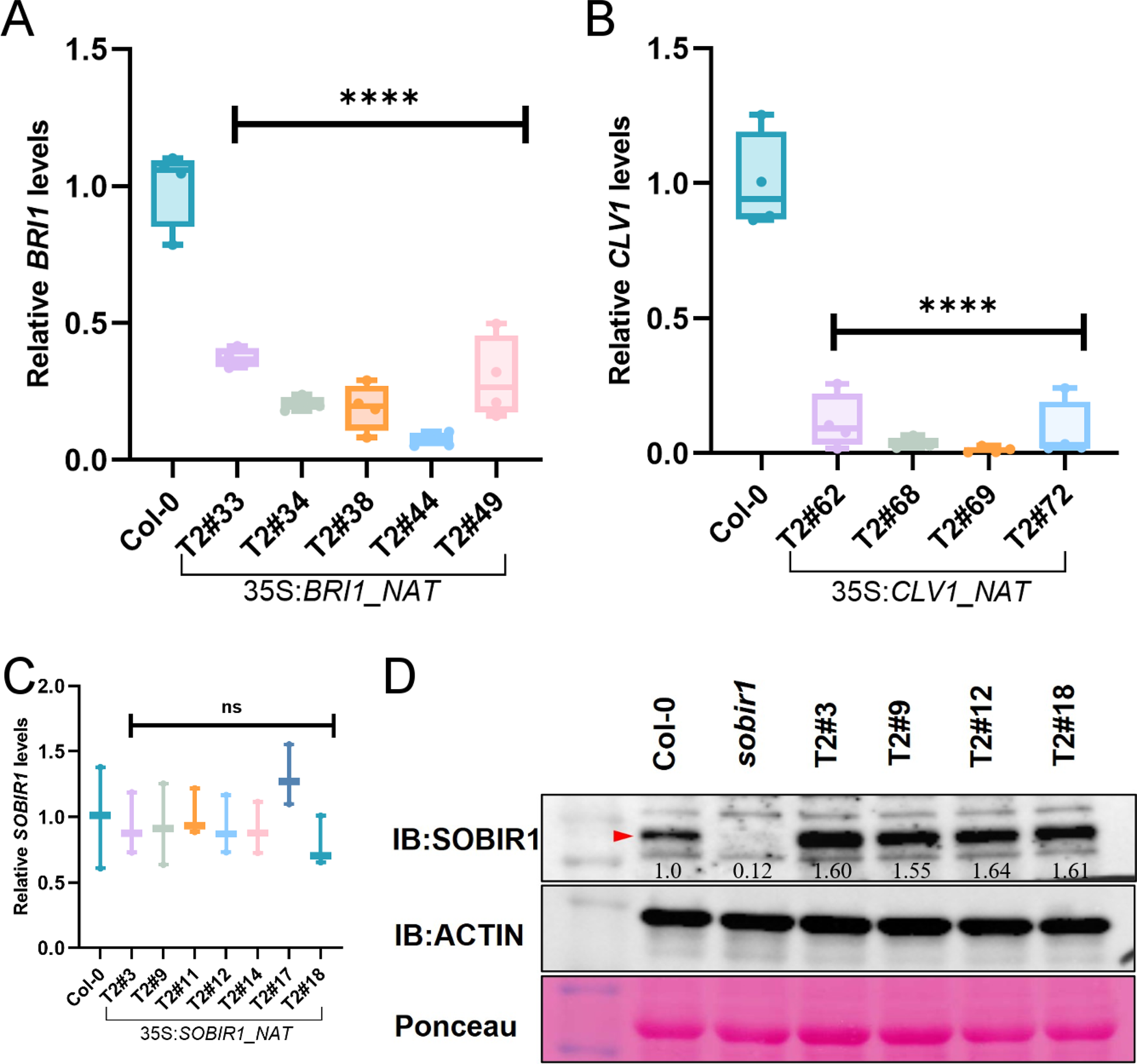
Overexpression of cis-NATs regulates LRR-RLK levels in transgenic lines. (A-C) Transcript levels of *BRI1*, *CLV1*, or *SOBIR1* LRR-RLK in transgenic lines overexpressing the corresponding NAT. For qPCR analyses, data was normalized using two internal controls – *Ubiquitin 2* and *PP2A*. For ease of visualization, the mean value of all Col-0 biological replicates was set to 1 and every data point was then calculated as fold-change relative to that value. Significance is determined by one-way ANOVA followed by Dunnett’s test for sample comparisons with Col-0 as control. Four biological samples and three technical replicates were analyzed, with the experiment repeated three times. (**** = p < 0.0001, ns= not significant (p > 0.05)). Closed circles represent the mean value for individual biological replicates, the boxes represent the first and third quartiles, the horizontal line indicates the median, and whiskers extend to the farthest points not considered outliers. (D) SOBIR1 protein levels in *SOBIR1_NAT* overexpression lines. 5-week-old plants of each genotype were infiltrated with OD 0.01 *Pst* DC3000 expressing the hopk1c effector and harvested after 24 hours. Relative quantification is shown inset. Each genotype sample was a pool of 4 different plants with 4 leaf discs harvested from each plant. The experiment was repeated three times with similar results.

As cis-NATs can also regulate their cognate gene through post-transcriptional and translational mechanisms(12, 21, 50), we then quantified the protein levels of SOBIR1 in *SOBIR1_NAT* overexpression lines. Basal SOBIR1 levels were below reliable detection, so we induced SOBIR1 protein accumulation in Col-0 and the *SOBIR1_NAT* overexpression lines by infiltrating with a strain of *Pst* DC3000 engineered to induce effector-triggered immunity (details in Materials and Methods). In induced tissue, *SOBIR1_NAT* overexpression lines consistently showed enhanced accumulation of SOBIR1 protein as compared to Col-0 plants, while the *sobir1* mutant showed no detectable SOBIR1 protein (Figure 3D). We also measured transcript levels of *SOBIR1_NAT* upon immune challenge in Col-0 plants and observed a significant upregulation in *SOBIR1_NAT* levels post-infection (Figure S3), suggesting that *SOBIR1_NAT* levels are highly responsive to immune elicitation. These results confirm that cis-NAT expression can regulate cognate LRR-RLKs using distinct mechanisms, consistent with previous reports(13, 15, 20, 50, 51).

### NATs display specific expression patterns *in planta*

To fine-tune LRR-RLK expression, cis-NATs must themselves be differentially expressed and we therefore assayed cis-NAT promoters for activity *in planta*. To visualize cis-NAT expression we created *β-glucuronidase (GUS)* reporter lines using the promoter of each cis-NAT and analysed expression and localization of the GUS signal in several independent transgenic lines (Figure 4A-L). All three tested promoters were active with varied expression intensities and distinct localization patterns. The *pBRI_NAT::GUS* lines showed the weakest signal, which was concentrated in the emerging leaves and petioles of mature plants. A distinct signal was visible in the leaf vasculature, while no visible staining was observed in the reproductive tissues and roots (Figure 4A-D). The *pCLV1_NAT::GUS* lines showed strong and broad expression in young cotyledons, roots, mature leaves, internodes of siliques, and flower sepals (Figure 4E-H). The *pSOBIR1_NAT::GUS* lines showed the most prominent localization in leaf vasculature (Figure 4I-L). These results show that the promoters of cis-NATs are active in plants with specific expression patterns consistent with a role in fine-tuning of LRR-RLK expression.

**Figure 4.**
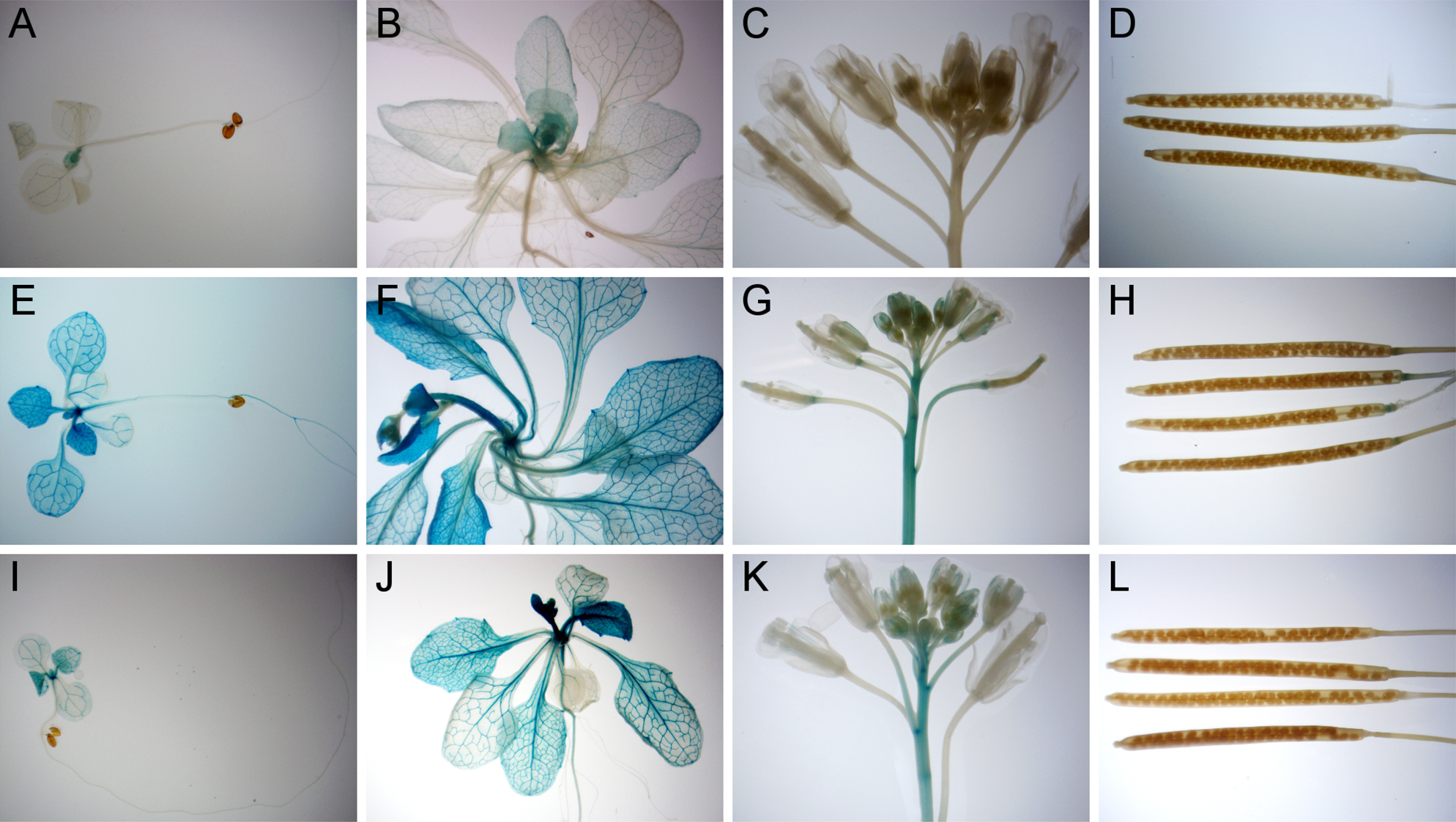
Expression pattern of cis-NATs. (A-D) Expression pattern of *BRI1_NAT* in seedlings (A), adult shoot (B), flowers (C) and siliques (D). (E-H) Expression pattern of *CLV1_NAT* in seedlings (E), adult shoot (F), flowers (G) and siliques (H). (I-L) Expression pattern of *SOBIR1_NAT* in seedlings (I), adult shoot (J), flowers (K) and siliques (L). (A-L) In all cases the seedlings used for GUS staining were 7-10 days post germination, adult shoots were from 3-week-old plants, siliques and flowers were from 5-week-old plants. At least 4 (up to 6) independent transgenic lines were stained for each cis-NAT, and all lines showed similar expression patterns.

### Tissue-specific expression of *BRI1_NAT* displays highly specific functional phenotypes

For cis-NAT regulation to produce the specific expression patterns observed in LRR-RLKs, the cis-NATs must function cell autonomously. To demonstrate that the functional effects of the cis-NATs are not mobile within the plant, we once again leveraged the *bri1* mutant system. The growth defects observed in *bri1* knockouts can be complemented by expression of *BRI1* in just the epidermal layer^30^. We therefore created transgenic lines that express *BRI1_NAT* under either an epidermis-specific promoter (*pAtML1*) or a sub-epidermis-specific promoter (*pPCAL*)(52).

Transgenic lines expressing *BRI1_NAT* under the epidermal promoter, in which we expect negative regulation of *BRI1* expression in the critical tissue, display the expected *bri1* mutant-like phenotypes in the T1 generation (Figure 5A). Of the 55 T1 plants, 26 displayed the *bri1* mutant-like phenotype. These plants also had significantly lower expression of *BRI1*, suggesting that *BRI1_NAT* can specifically reduce *BRI1* expression in the epidermal layer (Figure 5B). Transgenic lines expressing *BRI1_NAT* under the sub-epidermal promoter showed wild-type-like morphology. The 50 T1 plants displayed no aberrant phenotypes nor any signs of *BRI1* misregulation, which was confirmed by qPCR analysis of *BRI1* (Figure 5A, C). Sub-epidermal expression of *BRI1_NAT* had no regulatory or functional effect on *BRI1*. Critically, this suggests that the cis-NAT function is limited to the tissue layer in which it is expressed, allowing regulation of LRR-RLK expression in a cell autonomous manner.

**Figure 5.**
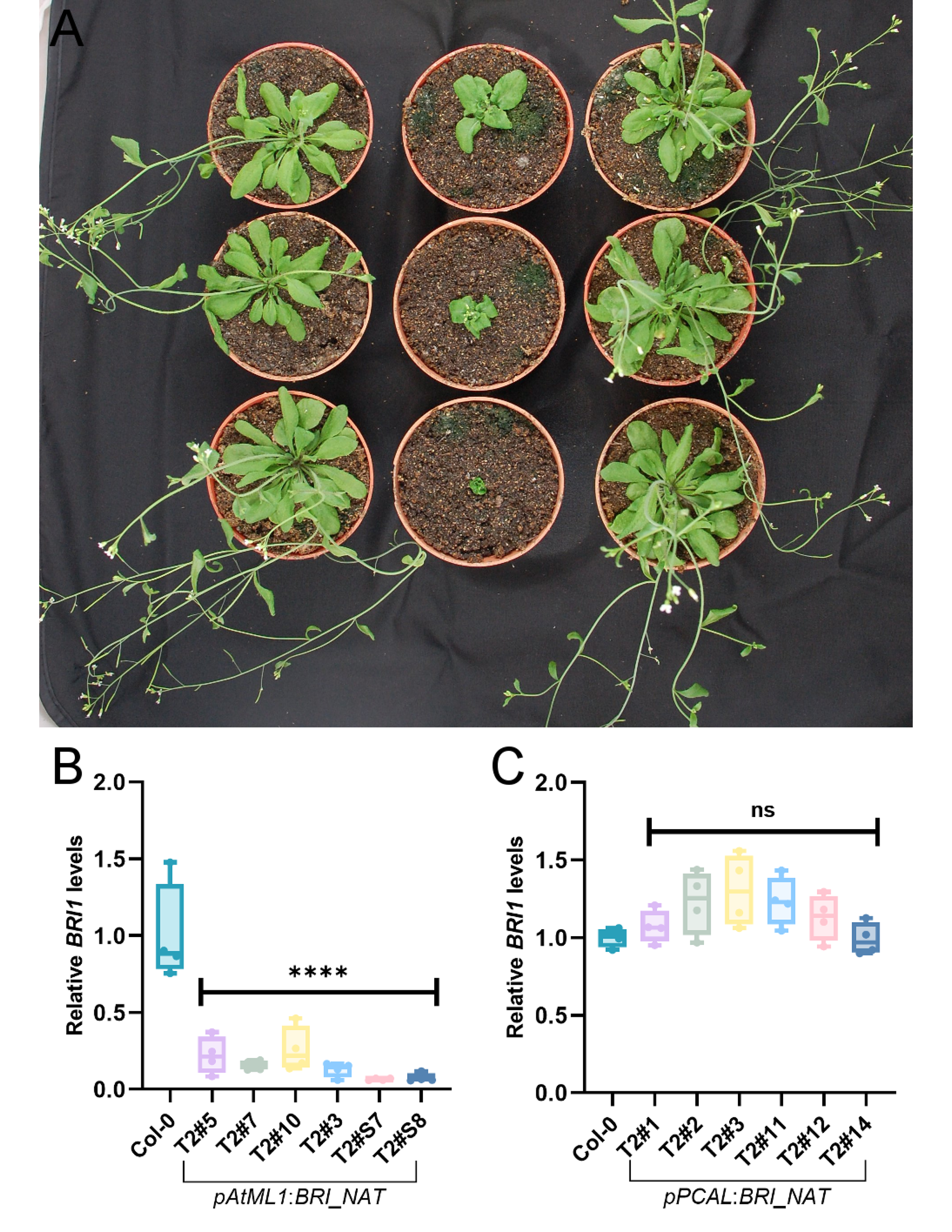
Tissue-specific expression of *BRI1_NAT* displays highly specific functional phenotypes. (A) Phenotypes of plants expressing *BRI1_NAT* under epidermal (*pAtML1*) and sub-epidermal (*pPCAL*) promoters. The left column contains 7-week-old Col-0 plants. The middle column contains 7-week-old transgenic plants expressing *BRI1_NAT* under the epidermal promoter *pAtML1*. The right column contains 7-week-old transgenic plants expressing *BRI1_NAT* under the sub-epidermal promoter *pPCAL*. Plants display a representative phenotype observed across several independent T1 lines (B-C) *BRI1* LRR-RLK mRNA levels in plants expressing *BRI1_NAT* under epidermal (*pAtML1*) promoter (B) or sub-epidermal (*pPCAL*) promoter (C). For qPCR analyses, data was normalized using two internal controls – *Ubiquitin 2* and *PP2A*. For ease of visualization, the mean value of all Col-0 biological replicates was set to 1 and every data point was then calculated as fold-change relative to that value. Significance is determined by one-way ANOVA followed by Dunnett’s test for sample comparisons with Col-0 as control. Four biological samples and three technical replicates were analyzed, with the experiment repeated three times. (**** = p < 0.0001, ns= not significant (p > 0.05)). Closed circles represent the mean value for individual biological replicates, the boxes represent the first and third quartiles, the horizontal line indicates the median, and whiskers extend to the farthest points not considered outliers.

### LRR-RLK genes are highly associated with cis-NATs across plant evolution

To test whether the association between LRR-RLK genes and cis-NATs was restricted to Arabidopsis, we analyzed publicly available strand-specific RNA sequencing data for multiple crop plant species. Accordingly, we compared our analyses to the number of Arabidopsis cis-NATs identified only in the Deforges et al. strand-specific sequencing dataset(12). Based on the breadth and depth of data available, we selected *Solanum lycopersicum* (tomato) as a dicot crop species and *Oryza sativa* (rice) as a more distantly related monocot species for study. In Arabidopsis, an associated cis-NAT was identified for 4273 of the 32833 total genes, of which 70 of the 225 LRR-RLK genes had an associated cis-NAT(12) (Figure 6). Overall, the LRR-RLKs are associated with cis-NATs 2.39 fold more than the genome on average, which is a highly significant increase in association (p = 1.09 x 10^−12^). Our analysis in tomato identified an associated cis-NAT for 5911 of the 35768 total genes, of which 54 of the 232 LRR-RLK genes had an associated cis-NAT (Figure 6). Tomato LRR-RLK genes are associated with cis-NATs 1.41 fold more than the genome on average, which is also a highly significant increase in association (p = 0.0048). Finally, in rice we identified an associated cis-NAT for 6303 of the 42189 total genes, of which 63 of the 316 LRR-RLK genes had an associated cis-NAT (Figure 6). Rice LRR-RLK genes are associated with cis-NATs 1.33 fold more than the genome on average, which is a highly significant increase in association (p = 0.0095). These results show that across a diverse group of dicot and monocot plants, LRR-RLK genes are much more likely to have an associated cis-NAT than genes in general. This conservation of association suggests a conserved and ancient role for cis-NATs in the fine-tuning of LRR-RLK expression in plants.

**Figure 6.**
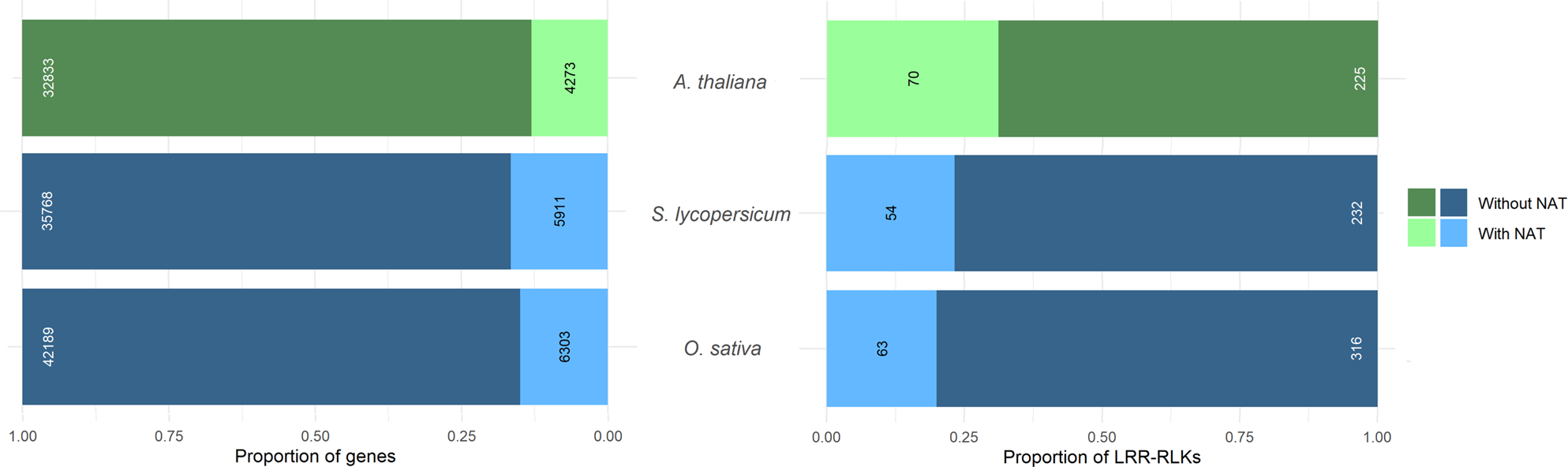
Association between LRR-RLKs and cis-NATs in *Arabidopsis thaliana*, *Solanum lycopersicum*, and *Oryza sativa*. For each species the total number of genes and cis-NATs identified are shown on the left. The number of LRR-RLK genes and associated cis-NATs are shown on the right. The total number of genes or LRR-RLKs are shown in white, while the number of cis-NATs in each set are shown in black. In all cases the bars show the proportion of genes with or without an associated cis-NAT, rather than absolute numbers. The data describing *A. thaliana* is taken from Deforges et al.(12) and shown in green for comparison with the analyses of tomato and rice shown in blue.

## Discussion

### The association of cis-NATs with LRR-RLK coding genes

In this study, we report a system-wide mechanism that plants employ to regulate the expression of LRR-RLKs via cis-NATs. By mining publicly available data, we found evidence that the vast majority of LRR-RLK genes in the Arabidopsis genome have an associated cis-NAT on the antisense strand (212/225, ∼94%). We confirmed that at least 115 of the 225 LRR-RLKs, or more than 50%, are associated with an actively expressed cis-NAT (Figure 1, Table S1) using an adaptor-based strand-specific qPCR approach in untreated 10-day old seedlings. cis-NAT expression is often restricted temporally, developmentally, spatially, or inducibly(7), suggesting that by testing only a single untreated plant stage we greatly underestimate the true extent of cis-NAT expression. A more expansive survey of expression in different tissues, developmental stages, and under different treatment conditions would likely confirm expression of many more predicted cis-NATs and would also show different expression levels for many confirmed cis-NATs. The near-universal association between LRR-RLK genes and cis-NATs stands in stark contrast to the genome-wide estimate of 15-20% of Arabidopsis loci reported to have an associated cis-NAT (7, 12). The discovery of a family-wide association between cis-NATs and the large LRR-RLK gene family raises the question of why this association exists.

It seems likely that the functions of the LRR-RLKs provide the first clue. LRR-RLKs are critical regulators of plant growth, development, and immunity. The expression of LRR-RLKs is therefore regulated developmentally, spatially, temporally, and inducibly to optimize plant fitness(31, 32). The misregulation of receptor expression can be catastrophic for the plant(53, 54), providing strong pressure for the development of a system capable of precise and robust expression control. Combinatorial control of gene expression by multiple transcription factors is known to allow the production of complex gene expression patterns from relatively few input features(55). Similarly, the additional layer of regulation provided by cis-NAT association would increase expression complexity beyond what is possible with transcription factors alone. Additional layers of regulatory control also increase the robustness of the system as a whole and enhance resilience to mutation and pathogen interference(56, 57). Cis-NAT regulation is uniquely specific, as each cis-NAT likely only regulates its cognate LRR-RLK, allowing for precise tuning of expression patterns in ways not possible through the activity of broadly acting transcription factors. We hypothesize that the requirement for precise control of LRR-RLK expression for plant fitness combined with the unique attributes of cis-NAT regulation has led to the conserved family-wide association.

### cis-NAT regulation of sense gene expression

The regulation of sense gene expression by associated cis-NATs can be positive or negative and can occur via numerous mechanisms(5, 7). The overexpression of *BRI1_NAT*, *CLV1_NAT*, and *SOBIR1_NAT* in Arabidopsis resulted in striking phenotypes, suggesting that cis-NAT dosage can modify sense gene function (Figure 2). Transcript levels of *BRI1* and *CLV1* were significantly downregulated in all cis-NAT overexpressing transgenic lines tested (Figure 3A, B), suggesting that the cis-NATs associated with these LRR-RLKs negatively regulate the sense gene. While *SOBIR1* transcript levels remained unchanged in *SOBIR1_NAT* overexpression lines (Figure 3C), SOBIR1 protein levels were significantly enhanced (Figure 3D), suggesting that *SOBIR1_NAT* acts positively at the translational level. The three characterized cis-NATs all regulate the activity of their cognate LRR-RLKs when overexpressed in trans, showing both phenotypic changes and regulation at the molecular level (Figure 2,3). Previous studies have also demonstrated that cis-NAT regulation can occur when the NAT is expressed in trans, which makes it possible to modulate the sense gene expression by overexpressing the NAT at a distant locus(12, 15, 19).

The known examples of cis-NAT-based sense gene regulation in the plant literature highlight the varied mechanisms through which they control agronomically important traits. These include negative regulation via histone modification(13), RNA polymerase II (RNAPII) collision(58), and modulation of mRNA stability(59). Other regulatory mechanisms have been reported in animal systems, including splicing regulation, nuclear retention or export of sense mRNA, and chromosome inactivation, but their existence in plants remains speculative(7, 8). These mechanisms have been shown to control critical agronomic traits in multiple plant species including flowering control, cold acclimation, biomass accumulation, phosphate homeostasis, and grain yield (13, 21, 50, 58–61). The discovery of the widespread association between the LRR-RLKs and cis-NATs presents a unique system to further explore possible regulatory mechanisms within plants and to determine whether there are cis-NAT features that can be used to predict regulatory mechanisms and function while potentially providing numerous opportunities to improve plant breeding.

### Specificity of cis-NAT expression

For cis-NATs to fine-tune the expression of their cognate LRR-RLK, they must display differential expression that overlaps at least in part with that of the LRR-RLK. The three cis-NATs tested showed distinct patterns of promoter activity, with at least partial overlap with known tissues where the cognate LRR-RLK is expressed(62–65) (Figure 4). Most described cis-NATs are either tissue-specific or induced by specific environmental or developmental signals(15). These specific and restricted expression patterns would allow cis-NATs to fine-tune the regulation of their cognate LRR-RLKs. As we move from this system-wide study towards a functional characterization of each cis-NAT, it will be intriguing to explore the spatio-temporal dynamics of this regulation at single-cell resolution. If cis-NATs are fine-tuning the expression of LRR-RLKs as we propose, then to maintain highly specific expression patterns the regulatory activity of the cis-NAT must be cell autonomous. By expressing the *BRI1_NAT* under tissue-specific promoters, we clearly show that its regulatory activity is restricted to the region where it is expressed (Figure 5). These features place cis-NATs in a position to fine-tune the expression of their cognate LRR-RLKs in a tissue-specific or inducible manner.

The negative regulation of *CLV1* and *BRI1* by their cognate cis-NATs produced the expected phenotypes mimicking those seen in LRR-RLK knock-out plants (Figures 2B and 2D). In the case of the positive regulation of SOBIR1 protein accumulation by its cognate cis-NAT, the resulting phenotype was not as expected. Previous reports have shown that strong overexpression of SOBIR1 results in constitutive immune activation, increased pathogen resistance, and widespread cell death, while moderate levels of overexpression did not result in an obvious phenotype(53). The *SOBIR1_NAT* overexpression lines accumulate SOBIR1 protein, display no noticeable cell death, and show increased susceptibility to infection with *Pst* DC3000 (Figure 2G). It is difficult to compare these studies directly as we only observe accumulation at the protein level, while the previous work showed overexpression at the transcript level by employing the strong, constitutively active 35S promoter. The lack of cell death may simply be due to dosage effects and not meeting the threshold level of SOBIR1 expression necessary to observe the reported phenotype, but the same reasoning would not explain the resulting bacterial susceptibility. One plausible explanation for this observation is that cis-NAT-dependent over-accumulation of SOBIR1 protein is spatially restricted to the cells or tissues where the SOBIR1 protein is normally present, since in the absence of *SOBIR1* transcript the cis-NAT will have nothing to act upon. In contrast to normal overexpression methods, this system results in increased SOBIR1 protein only in the context in which it is normally found and not in inappropriate tissues or throughout the plant. Such novel regulatory features make cis-NATs excellent tools to manipulate LRR-RLK levels in a natural context to gain a better understanding of their function. Cis-NATs that are positive regulators provide a unique tool to increase LRR-RLK levels at physiologically appropriate dosages without compromising the native localization of LRR-RLKs.

### Conservation of NAT association across plant evolution

Importantly, the observed association between LRR-RLKs and cis-NATs is not limited to the model organism Arabidopsis, which has an LRR-RLK repertoire that is not representative of all plants(66, 67). Highly significant enrichment of cis-NAT association with LRR-RLKs is observed in both the dicot *Solanum lycopersicum* (tomato) and the monocot *Oryza sativa* (rice). Given the diversity of these three plants, this strongly suggests that the association is evolutionarily ancient and conserved among plants. Due to the availability of data, this analysis was limited to only strand-specific RNA sequencing data from a limited number of samples and conditions. We have shown this identifies fewer overall cis-NATs than are truly expressed in Arabidopsis, and we therefore expect that as more crop species data becomes available we will likewise observe higher levels of cis-NAT association in tomato and rice.

This system-level study of the relationship between LRR-RLKs and cis-NATs is an important step toward understanding how the function of this critical receptor family is tightly controlled by a common mechanism to ensure optimal plant growth and survival. The regulation by cis-NATs provides an elegant example of the evolution of a shared regulatory mechanism to control the activity of a large family of LRR-RLKs with diverse functions, distinct expression patterns, and varied targets. This investigation of NAT-derived regulation of an entire gene family greatly expands on our limited understanding of cis-NAT regulatory mechanisms in plants(5, 7), and provides an ideal system to understand better the complex pathways through which cis-NATs regulate their cognate genes. Having demonstrated that this relationship between cis-NATs and LRR-RLKs is conserved across plant evolution also raises the possibility of engineering improved crop performance through cis-NAT modification, as they have been shown to control critical agronomic traits in other contexts(18, 21, 58, 59, 61).

## Material and Methods

### Screening for candidate cis-NATs and primer design

cis-NATs previously identified by strand-specific RNA sequencing, tiling arrays, and annotated in Araport11 were filtered for the ones associated with LRR-RLKs. Respectively, these sources identified 70 (Deforges et al. 2019)(12), 200 (Wang et al. 2014)(11), and 20 cis-NATs (Araport11)(40) associated with LRR-RLKs. Since many of the cis-NAT associated LRR-RLKs in this set were present in more than one dataset, we prioritized the datasets chosen for each cis-NAT detection. We prioritized the 70 cis-NATs associated with LRR-RLK genes from Deforges et al. as it was based on strand-specific sequencing. The Wang et al. predictions were used for all cis-NATs associated with LRR-RLK genes that were absent from Deforges et al. or that had an overlap with the Araport11 annotations. Araport11 had a very few annotated cis-NATs, so that source was only used for primer design if a particular cis-NAT was absent in the other sources. Based on these criteria, we narrowed down the total number of LRR-RLKs putatively associated with cis-NATs in our initial screening to 212 (Table S1). Of these, 70 cis-NATs were based on Deforges et al., 135 cis-NATs were from Wang et al., and 7 cis-NATs were from Araport11 (Table S1, Figure 1D). In cases where an LRR-RLK had more than one predicted cis-NAT from the same dataset, a single cis-NAT was chosen for primer design. Forward and reverse gene-specific qRT-PCR primers were designed in CLC Main Workbench (https://digitalinsights.qiagen.com) and are 20-25 base pairs in length with a melting temperature above 56°C. The 20-nucleotide adaptor sequence (GACTGGAGCACGAGGACACT) was added to the 5’ end of the reverse primer in each pair, which was used for reverse transcription as shown in Figure 1B. All primers used in this work can be found in Table S2.

### Plant material and growth conditions

*Arabidopsis thaliana* ecotype Columbia-0 (Col-0) was used as the wild type in this study. *clv-101* plants were kindly gifted by Prof. Zachary Nimchuk at the University of North Carolina at Chapel Hill. The T-DNA insertion mutant line for *SOBIR1* used in this study (GK-643F07, ABRC stock number CS461699) was purchased from ABRC and homozygous lines were isolated using insertion and gene-specific primers. Arabidopsis seeds used in the study were vapor-phase sterilized in a glass desiccator with 100 mL of bleach mixed with 3 mL of concentrated HCl to produce chlorine gas and treated for 3-4 hours, then stored at room temperature until needed. For the detection of cis-NATs, Arabidopsis seedlings were germinated on plates containing 1% agar, 0.5x Murashige and Skoog medium and 1% sucrose. Plated seeds were stratified at 4°C in the dark for 4 days to ensure uniform germination. Plates were then moved to a growth chamber (12 hours light at 120-150 µmol/m2s, 22°C during the day and 20°C at night, 55% relative humidity) and grown for 9-10 days. For the infection assays, sterilized seeds were germinated on soil and stratified in the dark at 4°C for 5-7 days before moving to the growth chamber. After germination, the seedlings were moved into individual pots and allowed to grow for 4-5 weeks until ready for the infection assay. Similar growth conditions were deployed for other experiments, and for harvesting seeds, plants were moved to long-day conditions after reaching maturity (16 hours light at 120-150 µmol/m2s, 22°C during the day and 20°C at night, 55% relative humidity).

### Transgenic lines construction

All overexpression lines for NATs were created using the binary vector PGBW602(68) and all promoter:GUS lines were created using the binary vector PMDC163(69). The corresponding overexpression or promoter fragment for each of the cis-NATs containing the attB sites was PCR amplified using Phusion polymerase (Thermo Scientific) and cloned into the gateway entry clone pDONR207 using BP clonase (Invitrogen) as per the manufacturer’s instructions. The fragments were subsequently introduced into the expression clones-PGWB602 or PMDC163 using LR clonase (Invitrogen) as per the manufacturer’s instructions. For cloning the *BRI1_NAT* (2146 bp) and *CLV1_NAT* (3448 bp) into the overexpression vector, sequences were directly retrieved from TAIR by using the coordinates identified in Deforges et al. For the *SOBIR1_NAT,* which is annotated as a 189bp NAT on TAIR, we used TSSPlant software (Softberry) to predict a putative transcription start site. Based on this prediction, we cloned a 304 bp fragment to overexpress *SOBIR1_NAT* in Arabidopsis. The promoter:GUS lines were created to contain the promoter regions upstream of the annotated/published start site of each cis-NAT (3kb upstream for *BRI1_NAT* and *CLV1_NAT*, 2kb upstream for *SOBIR_NAT*). For creating the tissue specific *BRI1_NAT* expressing lines, pCAL and pATML gateway vectors were used (52) The expression vectors were transformed into *Agrobacterium tumefaciens* (GV3101) and introduced into Col-0 plants by floral dipping method (70). The transgenic lines were screened using Basta or Hygromycin B.

### RNA extraction

Total RNA was extracted using RNazol RT reagent (Sigma-Aldrich) using the protocol described previously (71). Snap-frozen samples were crushed in liquid nitrogen and homogenized in 1 mL RNazol RT reagent followed by centrifugation at 15000 rcf at 4°C for 10 minutes. The supernatant was washed three times in chloroform to clear the aqueous phase. The samples were then mixed with isopropanol and incubated at −20°C overnight followed by centrifugation to pellet the RNA. The pellets were washed 3 times in 80% ethanol, then air-dried for 10 minutes. The collected RNA was resuspended in 30 µL of DEPC-treated water and stored at −80°C. RNA samples were treated with the DNase I kit (ThermoFisher) according to the manufacturer’s instructions prior to use.

### Quantitative PCR analysis

cDNA was synthesized using the M-MLV reverse transcriptase kit (Sigma-Aldrich) according to the manufacturer’s instructions. For cis-NAT detection, each cDNA reaction was comprised of 500 ng of RNA, the gene-specific reverse primer for the corresponding cis-NAT that contains the adaptor sequence, and the gene-specific reverse primers for the two internal controls, *Ubiquitin 2* and *Protein phosphatase 2a*.

For quantification of *BRI1* and *SOBIR1* genes, cDNA was synthesized from 1 ug of RNA using oligo-dT primers, as there was a long stretch of sequence in the coding region that did not overlap with the cis-NAT and could be used for sense gene detection using the oligo-dT-primed cDNA. For *CLV1* detection, cDNA was synthesized using a gene-specific reverse primer with an adaptor sequence, along with the gene-specific internal control primers.

For the cis-NAT and *CLV1* quantification, qPCR was performed using a gene-specific forward primer and an adaptor sequence containing reverse primer. The reverse primer was complementary to the adaptor sequence added during cDNA synthesis, ensuring that only the antisense strand was amplified (Figure 1C).

qPCR was performed using QuantStudio 3 Real-Time PCR system (ThermoFisher) with PowerUp SYBR Green Master Mix (ThermoFisher). For cis-NAT expression validation in seedlings, NATs were considered detected if a product with a single clean melt curve was amplified at a Ct value below 35 cycles and was detectable as a single distinct band on an agarose gel. qPCR data was analyzed using LinRegPCR software(72) using the process as outlined in the manual to calculate starting amounts of cDNA in each sample. To minimize any risk of biases arising from potential variation in reference gene expression levels, we employed two independent standard reference genes for the analysis (*Ubiquitin 2* and *Protein phosphatase 2a)*. Reference genes from each cDNA sample were co-amplified with the target to normalize RNA/cDNA content. The target gene abundances were initially determined relative to each of the reference genes, and the arithmetic mean of these two values was then used to calculate the overall relative target gene expression, that is represented in the graphs. For improved accessibility, we set the mean of datapoints for one condition in each graph as ‘1’ (i.e. wild-type condition) and compared the remaining mean expression values to this reference point.

### Bacterial infection assays

*Pseudomonas syringae* pv. tomato (Pst) DC3000 bacteria were grown overnight on King’s B media plates containing 50 µg/ml rifampicin at 28°C. Bacteria were resuspended in 10 mM MgSO_4_ to a final OD600 of 0.0005. Fully expanded Arabidopsis leaves (4-5 weeks old) grown under short day conditions were infiltrated with the bacterial suspension on the abaxial leaf surface using a 1-ml needleless syringe. Infected leaves were harvested (4 leaves per plant) on the third day post-infection for bacterial count. Each leaf was punched with a size 3 cork-borer and the harvested leaf discs (4 leaf discs from each plant, total surface area 1 cm^2^) were ground in 10mM MgSO_4_ solution after ethanol surface sterilization. For each genotype, 12 independent plants were harvested. Serial dilutions were plated on King’s B media plates and incubated for 2 days at 28°C, after which cfu was determined to measure the bacterial growth.

### Western blots

To quantify the SOBIR1 protein levels in the *SOBIR_NAT* overexpressing lines, SOBIR1 protein was induced by infiltrating the plants with PstDC3000 expressing Hopk1c (OD 0.01). After 24 hours, an equal amount of leaf tissue was harvested from the infected leaves and snap-frozen in liquid nitrogen. Each sample represented a pool of 4-5 independent plants, with 4 infected leaves from each plant. The collected leaf tissue was crushed to a fine powder and protein was extracted by homogenizing the tissue in GTEN buffer (10% glycerol, 25 mM Tris pH 7.5, 1 mM EDTA, 150 mM NaCl) with 10 mM DTT, 1% NP-40 and protease inhibitor cocktail (Complete, EDTA-free, Roche). The Supernatant was isolated and sample protein concentrations were calculated by Bradford assay (Bio-Rad Laboratories, 2010) using the Quick Start Bradford 1x Dye Reagent (Bio-Rad; Mississauga, CA). Samples were normalized to ensure gel loading of equivalent amounts. After normalization, extracts were denatured in Laemmli buffer by boiling at 95°C for 5 minutes. A total of 50 µg of protein from each sample was loaded on a 10% SDS-PAGE gel and protein electrophoresis was conducted using Bio-Rad Mini-PROTEAN Tetra System. Proteins were transferred from gels to the PVDF membrane using the Trans-Blot SD Semi-Dry Transfer Cell (Bio-Rad). Membranes were incubated in 5% nonfat dried milk in 1X TBST (Tris-buffered saline, 0.1% Tween 20) for 1 hour and immunoblotted with 1:1000 of anti-SOBIR1 antibody (Agrisera) overnight at 4°C in 2% milk solution in TBST. Following 4 washes with 1X TBST, membranes were incubated with 1:5000 dilution of anti-rabbit secondary antibody (Cedarlane; Burlington, CA) for 1 hour, then washed 4-5 times with TBST. Detection was performed using Bio-Rad ChemiDoc XRS+ System and Clarity Western luminol/enhancer solution (Bio-Rad; Missisauga, CA). Membranes were also stained with anti-Actin antibody (Agrisera) at a dilution of 1:5000 as a loading control, followed by incubation with an anti-mouse secondary antibody (1:5000, Agrisera). Ponceau S solution (VWR) was used to stain the PVDF membrane for an additional loading control. Quantification of bands was completed with the gel analyzer function as implemented in ImageJ 1.54g (73).

### GUS staining and imaging

Plants at various developmental stages were added to chilled 90% acetone solution, vacuum infiltrated for 10 minutes and incubated for 20 minutes at room temperature. The acetone solution was removed, and samples were washed several times with the buffer solution without X-gluc (0.2% Triton X-100, 50mM NaHPO4 Buffer (pH7.2), 2mM potassium ferrocyanide, 2mM potassium ferricyanide), and vacuum infiltrated in the last wash step for 10 minutes. The buffer solution was removed and replaced with buffer solution containing X-gluc at a final concentration of 2mM and vacuum infiltrated again for 10 minutes. Samples were incubated at 37°C for 12 hours and were subsequently washed with increasing concentration of ethanol (30%, 50%, 70% v/v) to remove chlorophyll. Samples were stored in 70% ethanol at 4°C until they were imaged. Samples were imaged on Optika Pro View stereoscope.

### Screening for NAT association in crop species

*Solanum lycopersicum* (ITAGv3.2) and *Oryza sativa subsp. japonica* (MSUv7.0) genomes and annotations were downloaded from Phytozome(74). Raw strand-specific RNA sequencing reads generated from the Illumina Tru-Seq stranded protocol were downloaded from the SRA (*S. lycopersicum* accessions SRX21480051 - SRX21480056, *O. sativa* accession SRX1528702). Reads were aligned to their respective genomes using HISAT2(75) with settings as default, except the dta-cufflinks option was enabled and the maximum intron length was set to the 99th percentile of intron lengths in the genome annotation. Transcripts were constructed using Cufflinks(76) with settings as default, except the minimum isoform fraction was reduced to 0.01 and the minimum frags per transfrag was reduced to 2. Putative cis-NATs complementary to a known gene were identified using Cuffcompare. Reads mapping to each assembled transcript were counted using HTseq(77) in count mode, and cis-NATs expressed at less than 1% of their sense gene expression level were discarded as potential false positives according to the strand specificity error rate of the sequencing kit. The coding potential of the remaining putative cis-NATs was assessed using FEELnc(78). The positive mRNA training set was generated using cDNA sequences from genes with evidence at the protein level according to UniProt annotations for each species(79). Where more than 5000 confirmed protein coding cDNA sequences were available, 5000 were selected randomly to train the classifier. The lncRNA training set was generated in shuffle mode and all other FEELnc parameters were set as default. LRR-RLKs were identified in each species according to Kileeg et al.(67). Briefly, protein sequences from each species containing an LRR domain identified by predict-phytoLRR(80), a kinase domain identified by hmmsearch(81), and lacking an NB-ARC domain were classified as LRR-RLKs. Hypergeometric tests were performed using phyper as implemented in the R programming environment version 4.3.1 (https://www.r-project.org/).

## Acknowledgments

We would like to thank Prof. Zachary Nimchuk at the University of North Carolina at Chapel Hill for providing the *clv101* seeds. We would like to thank Prof. Satyaki Rajavasireddy for providing feedback about drafts of this manuscript. This work was supported by the Natural Sciences and Engineering Research Council of Canada through a Discovery Grant Award to GAM and an Undergraduate Student Research Award to HK. HB was supported by a post-doctoral fellowship from the German Research Foundation (Deutsche Forschungsgemeinschaft).

## Supplementary figures

Figure S1. Melt curve and gel analysis of all the LRR-RLK associated cis-NATs detected in our study. The figure excludes *BRI1_NAT*, *CLV1_NAT,* and *SOBIR1_NAT* which are shown in Figure 1C. Only cis-NATs that had a clear melt-curve and gel band are shown as these were considered to be expressed NATs.

Figure S2. Genomic location of cis-NATs associated with *BRI1*, *CLV1 a*nd *SOBIR1*. Locations for *BRI1_NAT* and *CLV1_NAT* were identified from the study by Deforges et al. and the *SOBIR_NAT* location is based on Araport11 annotation. *BRI1_NAT* (2146 bp) and *CLV1_NAT* (3448 bp) display head-to-head overlap with their cognate LRR-RLKs with a majority of the NAT in the overlapping region. *SOBIR1_NAT* displays a full overlap. *SOBIR1_NAT* is annotated as a 189 bp NAT (cloned *SOBIR_NAT* region is 304 bp in length, based on transcription start size prediction, see materials and methods). Genes are shown across a 10 kb window in Jbrowse(82) with flanking gene(s) labelled. Black lines represent introns, blue bars represent UTRs, yellow bars represent exons, and red bars represent NATs. Arrows denote the 3’ end of the gene and direction of transcription.

Figure S3. *SOBIR1_NAT* levels 30 minutes post-infection with a strain of Pst DC3000 engineered to induce effector-triggered immunity. For qPCR analyses, data was normalized using two internal controls – *Ubiquitin 2* and *PP2A*. For ease of visualization, the mean value of all Col-0 biological replicates was set to 1 and every data point was then calculated as fold-change relative to that value. Significance is determined by one-way ANOVA followed by Dunnett’s test for sample comparisons with Col-0 as control. Six biological samples and three technical replicates were analyzed, with the experiment repeated three times. (* = p < 0.05). Closed circles represent the mean value for individual biological replicates, the boxes represent the first and third quartiles, the horizontal line indicates the median, and whiskers extend to the farthest points not considered outliers.

